# AutoPM3: Enhancing Variant Interpretation via LLM-driven PM3 Evidence Extraction from Scientific Literature

**DOI:** 10.1101/2024.10.29.621006

**Authors:** Shumin Li, Yiding Wang, Chi-Man Liu, Yuanhua Huang, Tak-Wah Lam, Ruibang Luo

**Author notes:** Those authors contributed equally (Shumin Li, Yiding Wang, Chi-Man Liu).

## Abstract

Rare diseases, affecting 300 million people globally, often result from genetic variants. Wholegenome sequencing has made variant detection more cost-effective, but interpreting these variants remains challenging. Current clinical practice combines quantitative evidence and literature, which is complex and time-consuming. We introduce AutoPM3, a method for automating the extraction of ACMG/AMP PM3 evidence from scientific literature using open-source LLMs. It combines an optimized RAG system for text comprehension and a TableLLM equipped with Text2SQL for data extraction. We evaluated AutoPM3 using our collected PM3-Bench, a dataset from ClinGen with 1,027 variant-publication pairs. AutoPM3 significantly outperformed other methods in variant hit and *in trans* variant identification, thanks to the four key modules. Additionally, we wrapped AutoPM3 with a user-friendly interface to enhance its accessibility. This study presents a powerful tool to improve rare disease diagnosis workflows by facilitating PM3-relevant evidence extraction from scientific literature.

## Introduction

Rare diseases remained a formidable public health challenge affecting ∼6% of the global population with ∼8000 known diseases ^1^. A key factor in diagnosing these rare diseases is the identification of the underlying genetic causes. While the rapid development of whole-genome sequencing (WGS) has made accurate variant detection more affordable, reaching a diagnosis for rare diseases can still be challenging, due to the intrinsic small sample size and limited current understanding of the complex relationships between genetic variants and their functional consequences.

The current clinical approach to variant classification relies on an evidence-based framework, primarily based on the 2015 guidelines published by the American College of Medical Genetics (ACMG) and the Association for Molecular Pathology (AMP) ^2^. As illustrated in Fig. 1A, the typical workflow for variant classification involves two major steps: variant annotation and literature review. Variant annotation often involves quantitative indicators, such as allele frequencies in populations (PP3), computational pathogenicity scores (PM2, e.g., REVEL ^3^), and the presence of the same change in known pathogenic variants (PS1). While variant annotation requires integrating various bioinformatics tools and databases, these well-defined evidence sources can be better handled through data-driven variant interpretation platforms, such as Exomiser ^4^, Genomiser ^5^ and Varsome ^6^.

**Figure 1.**
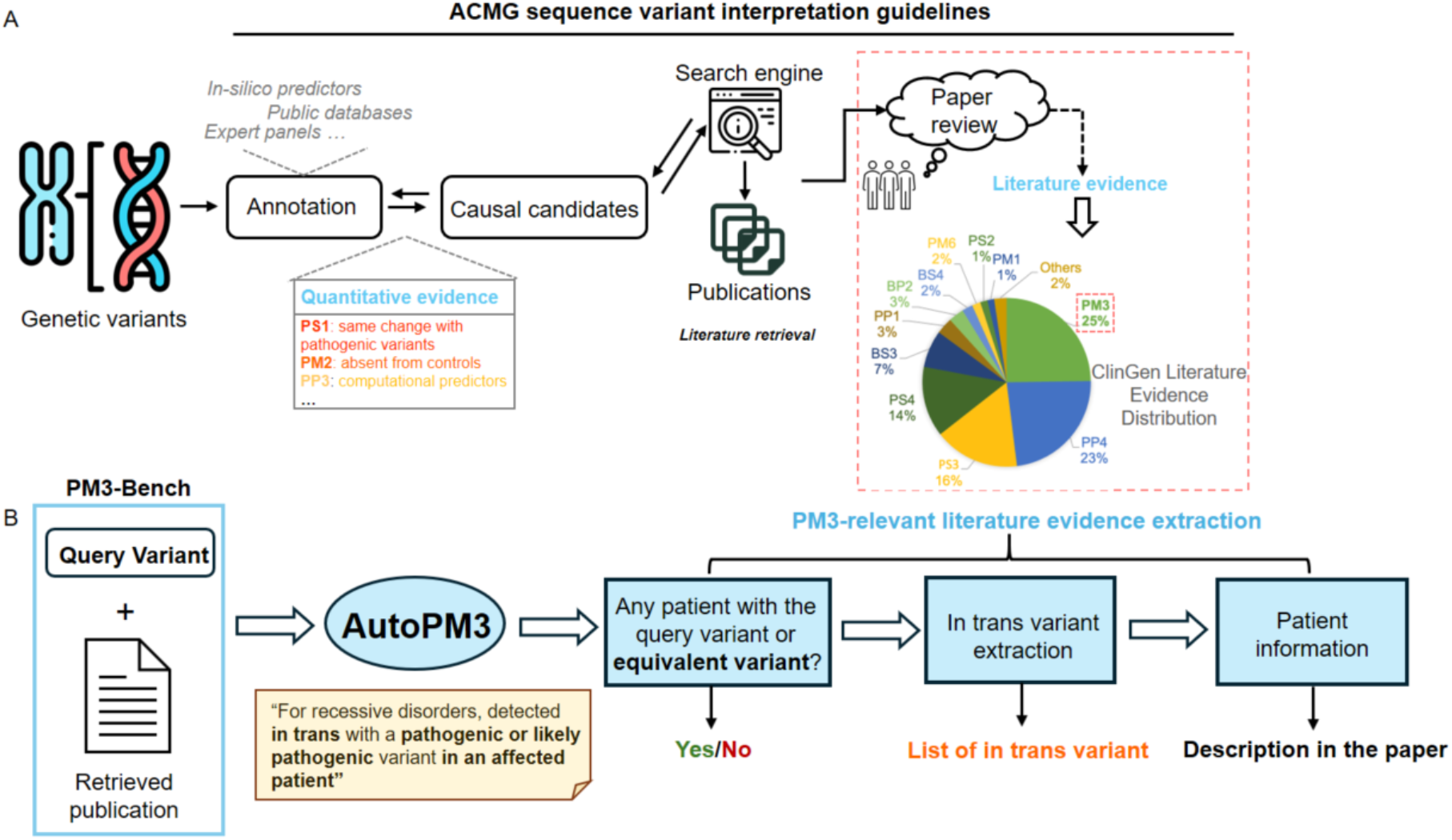
Overview of the variant interpretation workflow and the AutoPM3 system. A, the commonly adopted manual workflow for variant interpretation, involving variant annotation using external databases and tools, followed by manual literature evidence collection. B, the concept of the AutoPM3 system, which aims to automate the literature evidence collection step. Given a pair of a publication and a query genetic variant, AutoPM3 can extract relevant information from the literature, including whether the variant was reported, details about *in trans* variants, and potentially relevant patient information.

On the other hand, literature evidence represents another significant step of variant classification, with the fundamental aim of collecting known information from the whole scientific community. Literature evidence is broadly used in curated variants in ClinGen, with the PM3 criterion being the most frequently utilized, accounting for 25% of the literature evidence (Figure 1A). Specifically, the sequence variant interpretation (SVI) guidelines recommend a point-based system for determining the PM3 criterion, based on factors such as the number of probands, variant phasing, and the pathogenicity of the *in trans* variant ^7^. In this regard, a necessary step is to collect information on probands reported in the publications that share the same genomic variant and similar clinical syndromes with the target patient. Although biomedical entity-based literature retrieval tools, such as PubTator ^8^, have been developed, they lack the capability to fully understand the literature and extract the query-variant-oriented information and complex logical inferences required for variant classification. As a result, determining whether a publication contains supportive evidence is a time-consuming manual task that requires expensive expert curation.

Recently, large language models (LLM) have demonstrated remarkable capabilities in the biomedical field and become a promising option for understanding and extracting structured knowledge from publications or clinical notes ^9-11^. A recent work using a prompting strategy to demonstrate that GPT4-series models are potentially useful tools for identifying if one literature contains functional assay data and whether the data supports variant classification prioritization ^12^. Despite these investigations, the current applications of LLM-based data extractors either use in-context learning techniques or retrieval-augmented generation (RAG) frameworks, which are not optimized for the specific structures of scientific publications, particularly for tables, which are often the most information-intensive sections, nor are they designed for ACMG-criteria evidence extraction. Additionally, many of these LLM-based solutions rely on expensive GPT API services, which can limit their accessibility and prevent their use in local and offline environments.

To bridge this gap, we propose AutoPM3, a method for extracting PM3-relevant evidence from scientific literature, powered by open-sourced LLMs (Figure 1B). With the input of a query variant and a publication, AutoPM3 will determine if the publication mentions the variant, and identify *in trans* variants (if any). Specifically, the LLM-interpreted patient information is further aggregated to determine the PM3 criterion in the final output. The fundamental idea behind AutoPM3 is to separate the text and tables of the publication, and handle them using dedicated LLM-based modules. We employ text-to-SQL techniques to sensitively and accurately interpret the table data, while using an optimized retrieval-augmented generation (RAG) system to understand the main texts. There are four key modules in AutoPM3: variant augmentation, TableLLM, variant-specific retriever, and model fine-tuning. In addition, we have created a benchmarking dataset called PM3-Bench, based on ClinGen experts’ assertions comprising 1,027 publication-variant data points with ground truth of *in trans* variants and the number of patients.

Through benchmarking experiments, we have found that AutoPM3, equipped with lightweight models, significantly outperforms other methods in both variant hit and *in trans* variant identification, with an accuracy of 0.861 for variant hit and a recall rate of 72.5% for *in trans* variants. AutoPM3 also outperforms larger models, including LLaMA3:70B ^13^ and Mistral-Large:123B, under the systems of vanilla RAG or PaperQA ^14^. Through a sequential ablation study, we have demonstrated the effectiveness of AutoPM3’s key modules and how these modules can further increase the performance with larger models. In conclusion, AutoPM3 is an effective evidence extraction tool for PM3-relevant literature with small models and fast inference time, as well as lower hardware requirements. To make AutoPM3 accessible and easy to use for the community, we have wrapped it with a user-friendly interface using Streamlit, supporting both online and local usage.

## Results

### PM3-Bench: a comprehensive dataset for PM3 evidence extraction

Benchmarking datasets are crucial for developing and evaluating the performance of LLM-based systems. Although the ClinGen Evidence Repository provides expert-curated assertions, they are written in plain English, posing a difficult challenge for automated evaluation of benchmarks. To address this, we created PM3-Bench, a comprehensive dataset for PM3 literature evidence extraction, based on the ClinGen Evidence Repository (Figure 2).

**Figure 2.**
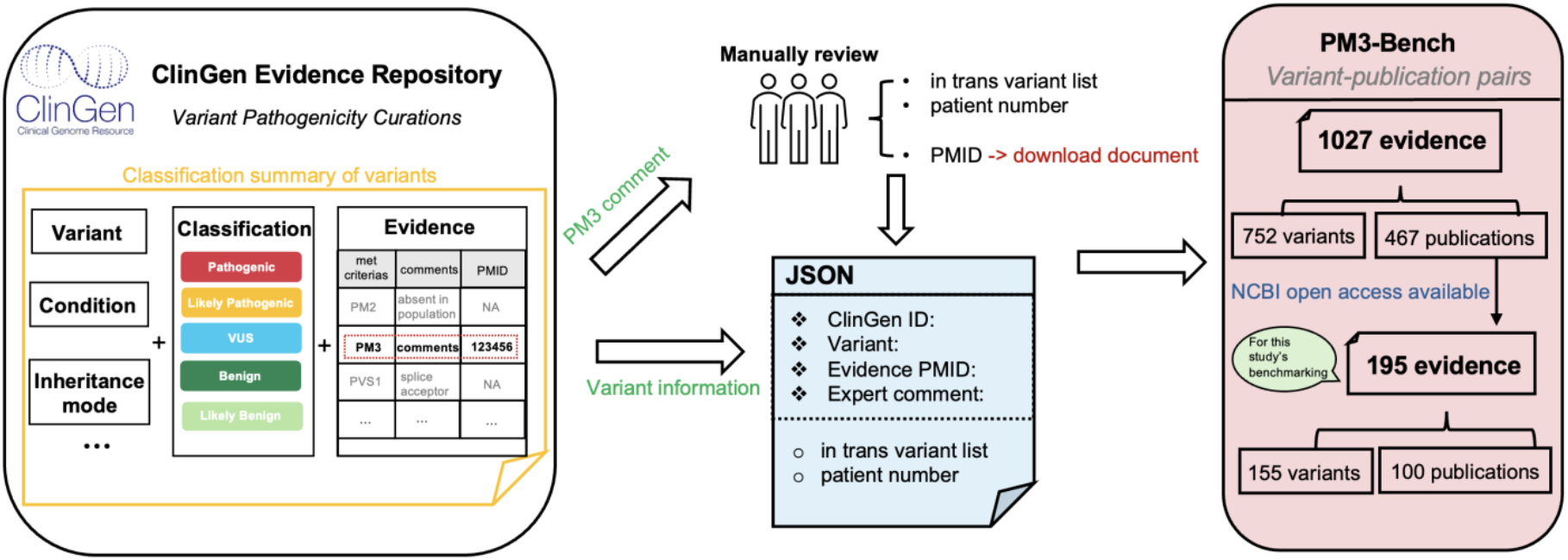
Data curation workflow for the PM3-Bench dataset. The process begins by collecting comments associated with PM3 from the ClinGen database. Comments that could be uniquely traced back to specific publications were retained. These variant-publication pairs were then formatted into a structured data format, and the comments were manually reviewed to extract key information into JSON data. In total, 1,027 evidence entries were obtained through this curation process. Of these, 195 entries could be downloaded from the NCBI Open Access subset and were designated as the testing samples for the performance evaluation.

First, we collected all variants with applied PM3 criteria from ClinGen. We then selected comments associated with PM3 if they were uniquely linked to a single publication, either through a traceable argument provided by ClinGen or if only one PMID was listed in the comment. This ensured that each comment and publication could be uniquely paired. For these comments, we manually extracted the *in trans* variants and the number of probands as the ground truth for the corresponding publication.

To evaluate the system’s capability to recognize ambiguous variant notations, we kept the target variants in the format of <*transcript identifier*>: **c**.<*position reference sequence*> <*alternate sequence***>** (e.g., NM_004004.5:c.71G>A). Additionally, we structured the extracted information and key details from ClinGen, including ClinGen ID, conditions, and PMID, into JSON format to facilitate further analysis.

The resulting PM3-Bench dataset includes 1,027 variant-publication evidence pairs, comprising 752 unique variants and 467 publications. Among these, publications that were available in the PMC Open Access Subset^15^ were downloaded, with 100 publications that were filtered by variants shown in supplementary files only, forming 195 variant-publication evidence pairs. We used these 195 samples as the testing data for this study, while the remaining 605 variant-comment pairs were utilized as fine-tuning samples (Methods). To facilitate the community’s use of PM3-Bench, we have released this dataset, aiming to provide open and reproducible benchmarks for PM3 evidence extraction tasks.

### Overview of AutoPM3

AutoPM3 is a document-level PM3 evidence extraction system powered by open-sourced LLMs. This system combines a TableLLM and a RAG module to separately process table and text content, involving four key modules (Figure 3).

**Figure 3.**
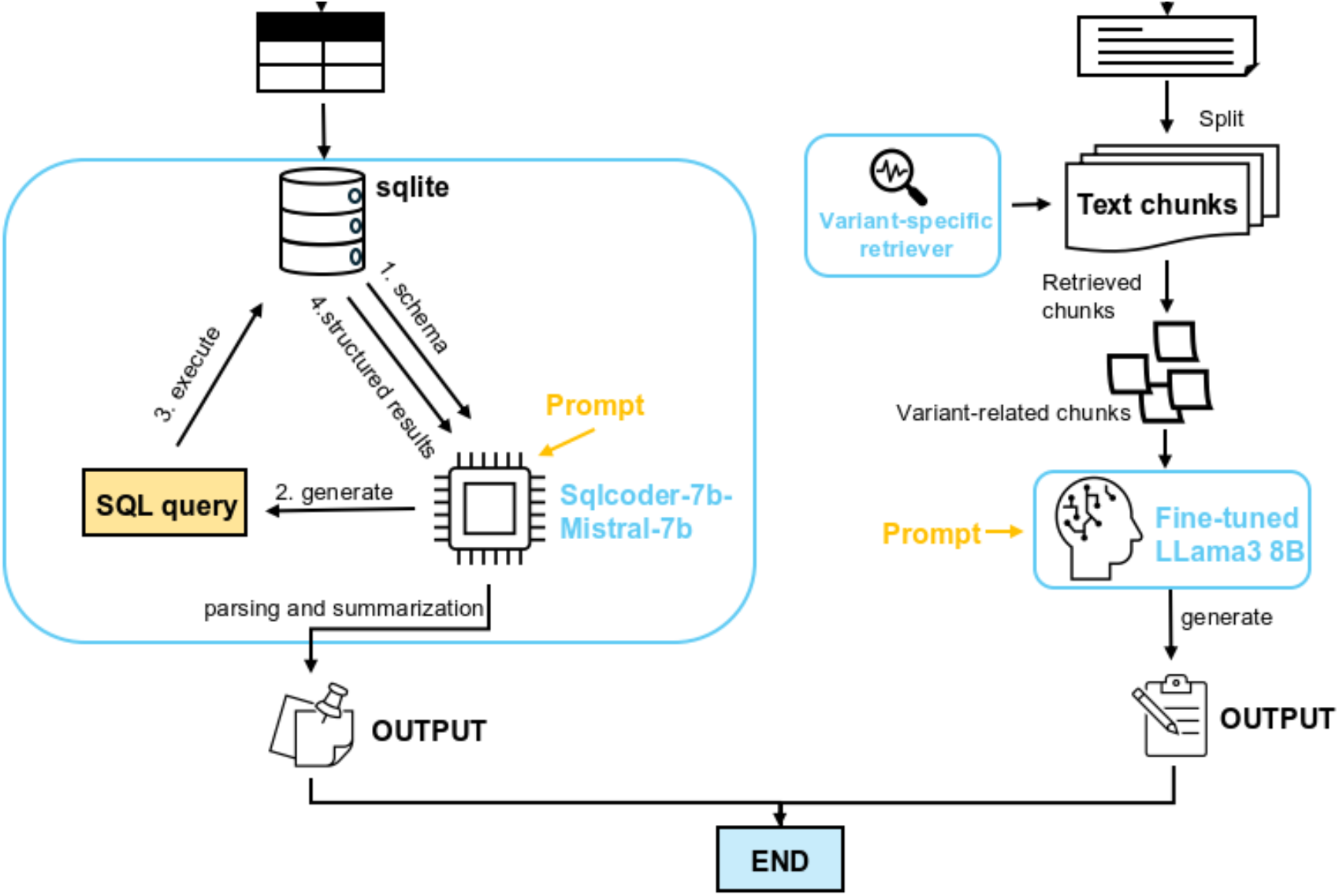
Overview of AutoPM3. The input to the system includes a literature file and a query genetic variant in HGVS format. The variant is first processed by a variant augmentation module, which expands the search space by generating different representations of the same variant. The literature file is then split into tables and main text content. The tables are stored in a SQLite database and processed by the TableLLM system, which utilizes a text-to-SQL core to generate and execute relevant queries and summarize the structured results using Sqlcoder-7B-Mistral-7B. The main text content is processed by an optimized RAG system. The variant retriever component is responsible for fetching variant-related text chunks to be used as context for the backend LLM, the fine-tuned LLaMA 3.8B model on default. The system used predefined queries for variant hit identification and in-trans variant detection, as well as support customized questions from the user.

First, given the query variant and the full content of a publication, we apply a variant augmentation module which uses the Mutalyzer API ^16^ to generate possible alternative representations of the variant, such as protein changes.

Next, our optimized RAG is utilized to extract PM3-relevant evidence for the query variant from the plain text. As in a typical RAG pipeline, the text is split into fixed-length chunks. Instead of using an existing retriever such as vector-based, we developed a variant-specific retriever to accurately identify chunks containing the query variant using regex matching which is sensitive to alternative representations of the variant (Methods). The retrieved variant-related chunks, along with the prompt, are then supplied to the LLM, for which we use a LoRA fine-tuned LlaMA3:8B as the default. AutoPM3 also supports larger open-sourced models like LlaMA3:70B and Mistral-Large:123B for evidence extraction (Methods).

For querying tables, which are often the most information-dense sections of scientific publications, we developed a TableLLM module. This module uses the sqlcoder-7b-Mistral-7B model to generate SQL queries based on the provided query variant and prompts, and then executes the generated queries against the SQLite database of the corresponding table (Methods). The SQL query results are then interpreted using the same model. Combined with the table header information, we parse and summarize the results into the final output.

We quantitatively evaluate AutoPM3 based on two key aspects. First, we assess whether the model can correctly identify the presence of the query variant in the publication. To evaluate potential hallucinations, we select an equal number of negative variants using real-world variants that were not reported in the publication. Performance is measured using sensitivity, specificity, and accuracy. Second, we examine whether the model can effectively detect the *in trans* variants based on the query variant, as their pathogenicity is vital for determining PM3. In this case, recall is used as the indicator. Notably, although not quantified, AutoPM3 can also generate summarized information about patients carrying the query variants, as presented in both tables and text.

### AutoPM3 effectively detects PM3-relevant evidence

Variant hit accuracy and *in trans* variant detection, which are the two vital factors for literature evidence extraction, were used for performance evaluation. For the variant hit task, we considered it a binary classification problem to determine whether a publication mentioned the query variant. In this experiment, AutoPM3 utilized two lightweight models: a fine-tuned Llama3:8B for RAG module and a 7B sqlcoder-mistral model for TableLLM. First, we compared AutoPM3 against the vanilla RAG framework, which employed the same prompts and chunk sizes but lacked AutoPM3’s four key modules. Instead, vanilla RAG utilized an ensemble retriever comprising BM25 and PubMedBERT embedding models ^17^ to ensure competitive retrieval performance ^18^. Additionally, we compared the performance of vanilla RAG equipped with models of different scales, including three open-source LLMs: Llama3:8B, Llama3:70B, and Mistral-large:123B. We also included PaperQA, a general scientific paper question-answering tool, with Llama3:70B as the backend, for comparison.

As shown in Figure 4A, AutoPM3 achieved the highest accuracy and F1 score of 0.861 and 0.865 for variant hit, significantly outperforming the second-best method, vanilla RAG (Llama3:8B), with an accuracy of 0.774 and F1 of 0.767. Notably, vanilla RAG and PaperQA using larger models (Llama3:70B and Mistral-Large:123B) exhibited variant hit accuracy of 10-26% lower than AutoPM3 equipped with small models (8B and 7B), despite being more than 9 times larger. However, these larger models achieved the highest specificity, all exceeding 0.96, partly by scarifying sensitivity (Supplementary Table 1). This discrepancy may result from the vanilla RAG retriever’s insensitivity to genetic variants, leading to more irrelevant text chunks retrieved and causing the larger models to have an inclination to answer negatively, resulting in high specificity and low sensitivity.

**Figure 4.**
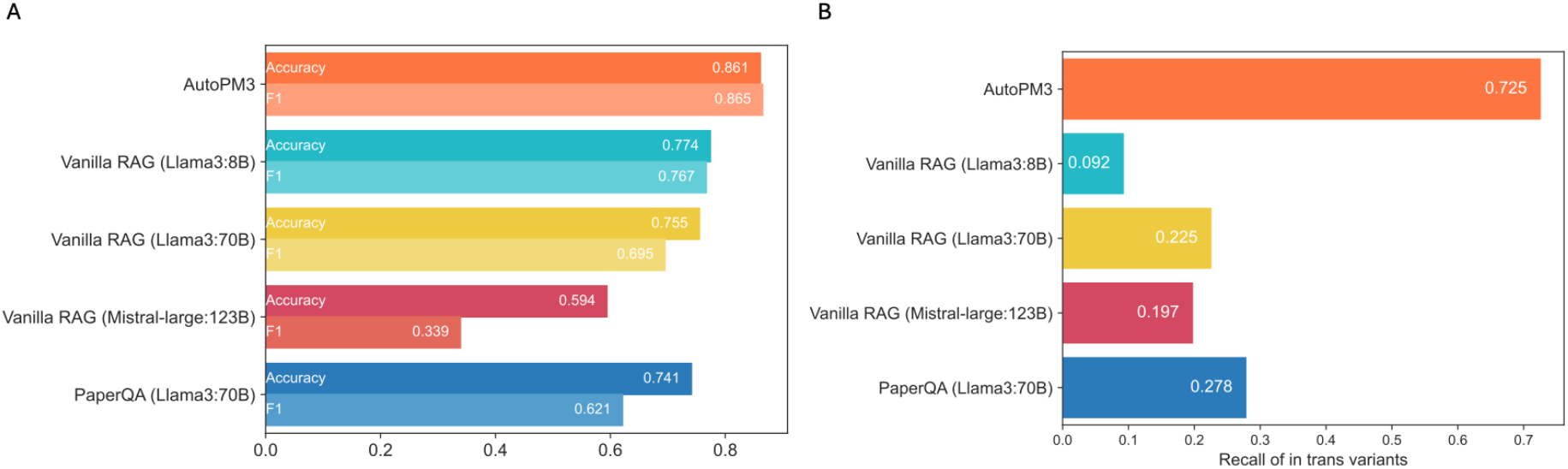
Benchmarking performance of AutoPM3 against baseline methods. (A) Comparison of variant hit identification performance across AutoPM3, vanilla RAG, and PaperQA. (B) Recall for *in trans* variant identification using the different methods. The input is the pair of positive query variant and literature, and the model generates the responses based on the predefined in trans specific query.

In addition to variant hit accuracy, we evaluated AutoPM3’s effectiveness in identifying the *in trans* variants of the patient. We examined if the ground truth in trans variant was summarized in the model’s answers and obtained the recall of *in trans* variant identification as the performance metric (see Methods). As demonstrated in Figure 4B, AutoPM3 identified 72.5% of *in trans* variants, while all other methods identified less than 30%, highlighting AutoPM3’s effectiveness, primarily benefiting from the four key modules. Of note, considering cumbersome detection of false hits, recall is the primary metric here for assessing in trans variants.

We observed that vanilla RAG (Llama3:8B), whose size can be considered the same as AutoPM3, identified only 9% of the ground truth *in trans* variants, failing in most cases. Moreover, even methods using large models like Llama3:70B and Mistral-large:123B, which were expected to exhibit better logical inference abilities, identified less than 30% of the ground truth *in trans* variants, which was substantially lower than AutoPM3’s recall, even though AutoPM3 was equipped with only 8B models. These results indicate that for complex inference tasks, such as *in trans* variant identification, large models may not exhibit superior performance compared to small models if the agent system is not optimized for table content and genetic variants, despite that large models usually outperform small models in general language tasks.

### Ablation analyses uncover key contributors of AutoPM3

Having demonstrated AutoPM3’s superiority over vanilla RAG and PaperQA in extracting PM3-relevant evidence, we further investigated the contribution of each key module: TableLLM, variant augmentation, variant retriever, and model fine-tuning. Moreover, we explored how large-scale models like Llama3:70B and Mistral-large:123B could further enhance the performance of AutoPM3 compared to the lightweight models for the RAG module.

We conducted experiments by sequentially adding these modules to three open-sourced models, starting with vanilla RAG (Supplementary Tables 2 and 3). For the variant hit task, shown in Figure 5A, variant augmentation resulted in minor accuracy increases for Llama3:70B and Mistral-Large:123B but decreased accuracy for Llama3:8B. This is because variant augmentation expands the search space, enhancing sensitivity but also increasing hallucination risks. While larger models maintained stable specificity, Llama3:8B’s specificity decreased from 0.8. to 0.692, with a 3% sensitivity increase. Furthermore, significant performance improvements were observed when adding the variant retriever to the Llama3:70B and Mistral-large:123B models. By retrieving only text chunks containing the query variant, the irrelevant chunks were minimized. Consequently, the sensitivity of Llama 3:70B and Mistral-Large:123B increased from 0.786 to 0.932 and 0.643 to 0.902, respectively. Additionally, Llama 3:8B’s specificity increased from 0.692 to 0.81. These results suggest the variant retriever module is crucial for 70B and 123B models to produce accurate answers and reduce hallucinations in 8B models. Next, we integrated the TableLLM with the RAG system of all models, dividing publications into text and table components. The text components were processed by the RAG system using variant augmentation and variant retriever modules, while the table components were handled by TableLLM. The performance of Llama 3:8B remained stable, while a slight fluctuation in accuracy was observed for Llama 3:70B and Mistral-large:123B. This suggests that for a simple binary task, large models already possess a certain level of table interpretation capability, reducing the additional benefit provided by TableLLM. Finally, we fine-tuned the Llama 3:8B model for evidence extraction, further increasing the accuracy of the 8B model-based system to 0.861.

**Figure 5.**
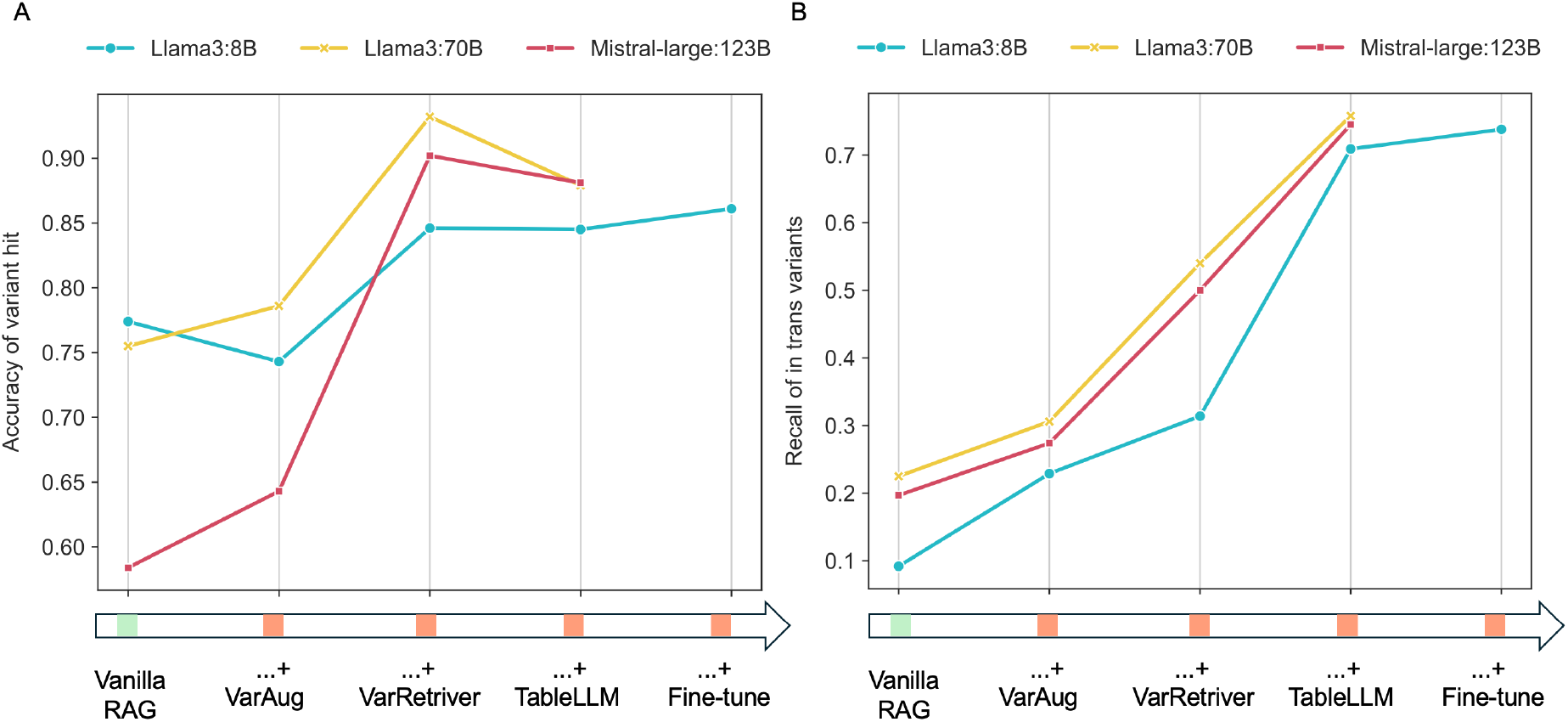
Contributions of AutoPM3 modules on PM3-relevant evidence extraction across different scaled language models. A, Variant hit identification accuracy when sequentially adding key modules of AutoPM3 to a baseline RAG system using LLaMA 3.8B, LLaMA 70B, and Mistral-Large 123B models. B, Recall of *in trans* variant identification when sequentially incorporating the AutoPM3 modules into the RAG systems with the different language model sizes.

For the *in trans* variant identification task, we observed a consistent performance increase when adding the variant augmentation and variant retriever modules for all models, as shown in Figure 5B. The variant retriever contributed the most significant performance jump, increasing recall by approximately 20% for 70B and 123B models and 10% for the 8B model. This also highlights the importance of relevant text chunks for comprehensive *in trans* variant detection. Notably, equipped with these two modules, both Llama 3:70B and Mistral-large:123B achieved over 50% recall, while Llama 3:8B remained around 31%, indicating a positive relationship between model size and *in trans* variant identification capability. We further integrated TableLLM into the system, consistently increasing the recall by approximately 20% for Llama 3:70B and Mistral-large:123B. The results indicated that although these two models demonstrated a certain level of logical inference for table content, their performance was inferior to the dedicated TableLLM module, necessitating its inclusion into the system. This effect was amplified for Llama 3:8B, whose table content comprehension capability was even weaker, resulting in a substantial recall increase by using TableLLM, from 0.314 to 0.709. In the final step, we fine-tuned the Llama 3:8B model to improve its understanding of genetics publications, boosting the overall *in trans* variant identification recall to approximately 0.725.

Through the sequential ablation study of our key modules on different model sizes for PM3-relevant evidence extraction, we observed performance gains driven by our key modules. Specifically, the variant retriever benefited the Llama 3:70B and Mistral-Large:123B models the most, primarily because their general capability was limited by low-quality retrieved context, an issue addressed by the variant retriever. Another vital module was TableLLM. We noticed that Llama 3:8B had limited capability in understanding table content, especially for tasks requiring logical inference like in trans variant identification. In this scenario, we adopted the lightweight TableLLM for table content through the text2SQL approach, effectively addressing this issue. On the other hand, TableLLM, with the size of 7B, outperformed larger models, namely Llama 3:70B and Mistral-Large:123B, when handling table content. Therefore, in our lightweight version, we used fine-tuned Llama 3 for text and TableLLM for tables, balancing time cost and accuracy, while it should be noticed that AutoPM3 also supports larger models such as Llama3: 70B and Mistral-Large: 123B for the RAG module, which has better capability than the default Llama3:8B model but requires much more computing resources.

### Demonstrating the utility of AutoPM3 through use cases

To allow the quantitative analysis capabilities of AutoPM3 to be exploited more easily, we have integrated AutoPM3 with an intuitive browser-based interface known as Streamlit. This interface can be accessed online at https://www.bio8.cs.hku.hk/autopm3-demo/ or deployed locally. The public demo version utilizes AutoPM3 equipped with the RAG modules of fine-tuned Llama3:8B, while users have the option to switch to larger models in their local deployment.

As illustrated in Figure 6A, the system requires two inputs: the query variant in HGVS notation and the PMID of the publication of interest. The system retrieves the corresponding publication using the NCBI Open Access API and then executes AutoPM3 in the server backend. Upon completion, the results are displayed on the same page (Figure 6B).

**Figure 6.**
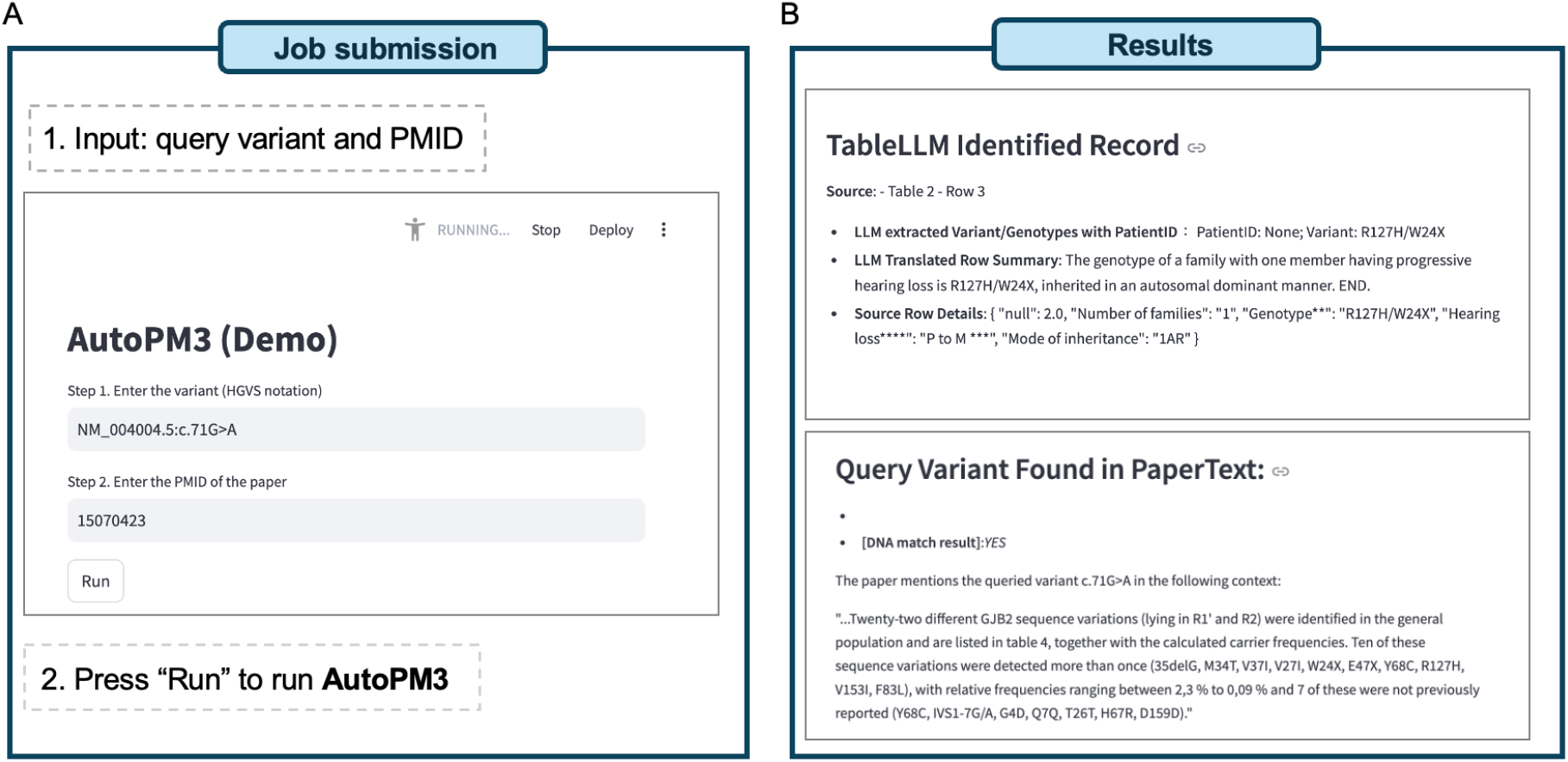
Overview of the AutoPM3 Interface and Use Cases. A, Illustration of the input and usage of AutoPM3. B, Example outputs generated by AutoPM3, showing results from both TableLLM and the RAG module.

The results are divided into two sections based on TableLLM and RAG modules. For TableLLM, each queried result is presented in four lines of information: 1, The source index of the results, e.g., “Table 1, row 6”; 2, inferred genotypes per patient; 3, summarized information of identified rows; 4, the original row in the table. It ensures that PM3-relevant information extracted from tables can be easily traced back to the original publication manually. Similarly, for the RAG module, which processes text, the rationale for the generated content follows the same structure. The results are presented based on two predefined queries: variant hit and *in trans* variant detection. Additionally, the results accommodate different formats of variants, including DNA change and protein change formats. The context of the query variant in the original text is extracted by the LLM for comprehensive consideration.

## Discussion

In this study, we present AutoPM3, a framework that utilizes LLMs for the efficient extraction of PM3-relevant evidence from the literature. Alongside this framework, we compiled PM3-Bench, a new dataset comprising variant publication pairs with curated ground truth, designed to facilitate both development and evaluation within the research community. Through the integration of AutoPM3 and PM3-Bench, we explore the potential and effectiveness of an LLM-based approach to enhance the document-level literature evidence collection process for the ACMG-guided variant interpretation workflow.

The core of AutoPM3 involves integrating two distinct modules for independently processing tables and text. For table-based evidence extraction, we framed the task as a database query problem and developed the TableLLM module. This component automatically generates SQL executable commands using the sqlcoder-Mistral-7B model, enabling database operations on tabular data. The results retrieved from the tables are then translated into summarization texts using the same LLM, for the determination of PM3-relevant evidence. Concurrently, we employed the RAG pipeline to extract evidence from the main text of publications. We optimized the vanilla RAG by developing a variant retriever and fine-tuning the backend LLM, Llama3:8B, for our experiments. The combination of TableLLM and RAG within AutoPM3 demonstrates effective document-level extraction, significantly outperforming both vanilla RAG and PaperQA, even when these systems were powered by models 9 times larger. The design of AutoPM3 powered by two lightweight models, ensures fast response times and lower hardware requirements for local deployment.

Furthermore, we conducted a sequential ablation analysis of AutoPM3’s key modules, exploring performance enhancements for the RAG module through the utilization of larger models. Our analysis revealed that the variant retriever integrated within the RAG system is crucial for enhancing performance in both variant hit and *in trans* variant identification. By employing a sensitive matching strategy, the variant retriever effectively locates text chunks containing the query variant, unlocking the potential of larger models (70B and 123B) and achieving approximately a 20% increase in performance. TableLLM also plays a vital role, particularly in *in trans* variant identification. While RAG with the variant retriever successfully retrieves relevant text chunks, challenges remain in accurately understanding tabular data. Although larger models perform better, their recall of *in trans* variants is still capped at around 0.5. In contrast, our TableLLM reformulates the task as a database query problem, effectively addressing this limitation and improving recall by over 20% when integrated with RAG across all model sizes. While the RAG system of AutoPM3 can be further improved by using larger models as the backend, we did not include the 70B and 123B models in our default settings due to their increased running time and hardware requirements. However, the inherent support for these models is maintained in the AutoPM3.

In addition to the quantitative indicators of variant hit and *in trans* variant, the output of AutoPM3 also summarizes the patient information associated with the query variant described in the document. Our principle is to extract the PM3-relevant evidence from the literature, instead of directly classifying whether a paper satisfies PM3 or not, as this approach is more robust, allowing users to review the results and corresponding source texts to make the final decision. In this regard, for future work, we will investigate the optimal interactive way between AI-based applications like AutoPM3 and human experts to minimize costs and risks. Another direction is to explore linking external pathogenicity databases to AutoPM3, which can further enrich the PM3-relevant information by justifying the pathogenicity of identified *in trans* variants.

In addition, one limitation we observed is the file format of the publications. Although we used NCBI-provided XML files in our experiments, the most common format of publications is PDF. While AutoPM3 is applicable to PDF files, its performance partly depends on the accuracy of the PDF parser used for extracting tables. With the development of more accurate PDF parsers, this issue can be addressed in the near future.

In summary, this work demonstrates that utilizing the AutoPM3 framework, open-sourced LLM-based literature evidence collection for variant interpretation is a promising and effective approach. With lightweight models and a user-friendly interface, AutoPM3 can be easily installed and run locally within a clinical center, with minimum costs. For instance, using an Nvidia RTX 4090, it averages 74 seconds per variant-publication pair in our evaluation samples. Furthermore, we have made our PM3-Bench dataset available to the research community, allowing for the investigation of LLM-based methods with ground truth datasets. We expect these advancements to facilitate the literature evidence collection for the variant interpretation workflow, thus supporting rare disease diagnosis.

## Methods

The AutoPM3 system consists of two main components: TableLLM and RAG. Prior to these two modules, we implemented a variant augmentation step to recognize potentially different formats of the same genetic variant. The implementations of these modules are described in the following sections.

### Variant augmentation

Genetic variants can be represented using either DNA or protein change notations in the HGVS format. To standardize these descriptions, we utilized the Mutalyzer 3 normalization API for variant augmentation ^16^. When querying with an HGVS notation, our system searches for both the DNA and protein change formats as independent inputs to the downstream TableLLM and RAG modules.

### TableLLM

We developed the TableLLM module with the core of a Text-to-SQL component, aiming to enable structured data retrieval and understanding, addressing the limitations of current open-sourced LLMs. Tables were extracted from the input publication using BioC ^19^, and a lightweight database (SQLite3) was established for storing each structured table. The processing involves three main steps: 1, SQL commands generation, 2, SQL execution and results fetching, and 3, post-processing and convertion to texts. Text2SQL generates the SQL commands by utilizing useful information from the database, including table headers and schema, as part of the prompt for the specific LLM, sqlcoder-7b-Mistral-7B-Instruct-v0.2-slerp (see Supplementary Information). To achieve optimal precision and recall in queries for variant hits and *in trans* variant detection, we implemented two measures: Since gene symbols can vary according to different standards, the generated SQL is guided by matching variant positions only, followed by subsequent filtering on the retrieved records. Specifically, the filtering is implemented by pattern-matching between fetched string and variant position. The pattern detects if a previously matched position string appears only among statistical digits to remove false positive records, like “frequency is 0.93884” in certain records with query position at “388”. For documents containing more than five tables, tables are split into chunks for recursive Text2SQL processing to avoid performance degradation as the input context of the LLM grows. Specifically, for *in trans* variant detection, TableLLM will first identify the rows of the query variant and then get its potential ID by the first column. The upstream and downstream rows with the same ID will also be extracted for further processing. Lastly, we applied the same LLM to the fetched records with specific prompts (see Supplementary Information) to generate user-friendly outputs. These outputs interpret table records back into plain text, highlighting key elements such as patient information and genotype. Moreover, a source-tracing feature indicating table and row numbers is provided, enabling users to review the original documents if necessary.

### The RAG module

AutoPM3 adopts the RAG pipeline for evidence extraction based on main texts of the literature. The texts were first divided into text chunks with 1500 characters size and each chunk has 100 characters overlap, RecursiveCharacterTextSplitter of LangChain was used for this purpose. Given a query variant, we applied the variant retriever to these chunks and retrieved at most 5 chunks containing the variants as the context. The prompt and queried question for variant hit and *in trans* variant detection were listed in Supplementary Info. QA chain of LangChain with chain type set as “stuff” was used for answer generations. The backbone LLM of the RAG pipeline was locally hosted by Ollama, using either our fine-tuned Llama3:8B or other open-sourced LLMs, including Llama3:8B, Llama3:70B, Mistral-Large:123B, etc.

### Variant-specific retriever

Retrieving relevant document chunks is crucial for a RAG pipeline. For PM3-relevant evidence extraction, a relevant chunk almost certainly mentions the query variant in some representation. Unfortunately, typical retrieval methods such as vector databases cannot handle variant notations well enough. This is because variants are usually represented as a mix of digits, letters and symbols without word boundaries, thus confusing the tokenizer. To improve the chance of retrieving relevant chunks, we have created our own retriever which finds the chunks explicitly mentioning the query variant.

As in variant augmentation, the query variant is first converted to different representations using Mutalyzer 3. Examples of representations include DNA change (c.274G>T), protein change (Asp92Tyr), and its shorthand (D92Y). To increase the chance of matching the variant in the text, we create regular expressions from the different representations, e.g., allowing whitespaces, omitting parentheses, interchanging X and the asterisk (*), etc. Successfully matched chunks will be retrieved for downstream query. In case there are too few matched chunks, we will relax the condition to matching digits only (e.g. matching “274” in “c.274G>T”), or even allow some error in the numeric value (e.g. 272-276) since the variant position may not be identical across reference sequences.

### Model fine-tuning

Common 8B LLMs tend to generate lengthy responses through explicit limitation is given in the prompt, bringing up hallucination and difficulty in reading and evaluating the correctness of responses. Thus, succinct answers are preferred to only respond user’s query with clear evidence of PM3 relevant information such as genotype, patient and parental info. Fine-tuning is feasible to meet this objective with affordable GPU memory size and training efficiency provided by LoRA fine-tuning ^20^ on LLama3:8B via Pytorch and Deepspeed training-time acceleration ^21^. With LoRA, parameters from only each transformer layer are modulated by an additional low rank layer. Statistically, low rank matrices only introduce 54,525,952 trainable parameters compared to original LLama3:8B with 8,084,787,200 parameters totally, accounting for 0.674% only. The trainset is extracted from curator reports from ClinGen with relevant comments as inputs and *in trans* variants as true labels, all testing samples were excluded. In this way, the model is trained to further learn representations of gene symbol and variant abbreviation and generate only short and clear answers for target variant and relevant *in trans* QA. The example of a training sample is shown in Supplementary Information. We then fine-tuned the Llama3:8B on 3 local-rank GPU nodes by torch-distributed-parallel along with Deepspeed at zero3 level. The parameters are as follows: *num_train_epochs* was set at 100, *gradient_accumulation_steps* is set at 8, *learning_rate* is set at 1e-5, *weight_decay* is set at 0.1, *adam beta2* is set at 0.95, *warmup_ratio* is set at 0.01, *lr_scheduler_type* is set at cosine and *model_max_length* is confined to 4096.

### Related methods

#### PaperQA

PaperQA is an LLM-based tool for doing high-accuracy RAG on scientific publications. It excels at information retrieval, summarization, and citation. Although PaperQA is not fine-tuned for genetics and medical literature, its superior performance in understanding scientific knowledge makes it an excellent tool to compare with. We benchmarked PaperQA version 4.9.10 on our PM3-Bench dataset. Since PaperQA required following a lot of instructions, we chose Llama3:70B as the underlying model. It is worth noting that PaperQA was designed to generate answers in well-written paragraphs, and we failed to force the output into simpler formats such as a simple Yes/No or a list of variants, by prompting. As such, we decided to evaluate the performance of PaperQA by manually inspecting its outputs for all test cases.

#### Vanilla RAG

We implemented a vanilla RAG framework using LangChain, similar to the RAG module employed in AutoPM3. The full text of the publication was split into text chunks based on characters, with a chunk size of 1500 and an overlap of 100 characters. The text chunks were indexed using a PubMedBERT embedding model and stored in a FAISS-based retrieval system. The retrieval process utilized an ensemble approach, combining an embedding-based retriever and a BM25 retriever, with equal weights of 0.5 assigned to each. For both retrieval methods, the top 2 most relevant chunks were selected. The prompt and query questions used in the vanilla RAG framework were the same as those employed in the RAG module of AutoPM3, as described above. We utilized the QA chain in LangChain, with the chain_type set to “stuff” and the utilized LLMs were locally hosted and downloaded by Ollama ^22^.

### Evaluation

#### Variant hit

We evaluated the variant hit task as a binary classification problem, where the goal was to determine whether a publication mentioned the queried genetic variant. The positive samples were obtained from the PM3-Bench dataset, which were collected from ClinGen. To generate the negative samples, we randomly selected variants mentioned in a different publication and located in a different gene from the true variant set. We then performed regular expression matching to ensure that these negative variants were not mentioned in the corresponding publication. The final testing dataset consisted of 195 positive pairs and 195 negative pairs.

For the evaluation, we used the following approaches: for TableLLM, if TableLLM reported any queried results, we considered it as a positive answer; otherwise, it was treated as a negative answer. For RAG, the system generated responses based on the question “*Does the context mention the queried variant {variant name} and what is the surrounding context? if such variant exists, say *YES* at first otherwise say *None* (focus on variant: {variant name})*”. If the response contained the word “Yes”, we considered it as a positive answer; otherwise, it was treated as a negative answer. If either TableLLM or RAG module reports a positive response, we consider it as the positive answer. The accuracy, sensitivity and specificity, precision and F1 of variant hit were calculated as below (note, the numbers of positive and negative samples are equal):

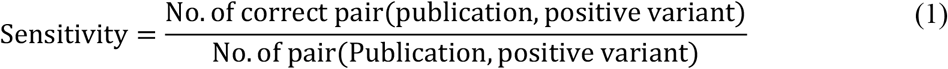

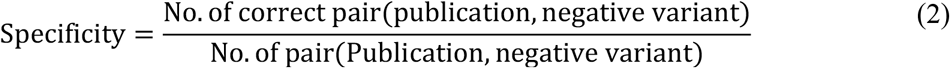

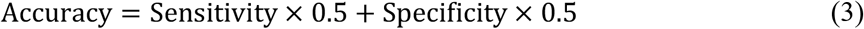

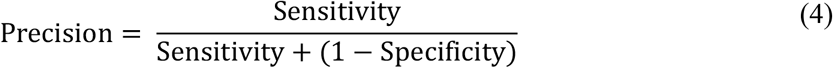

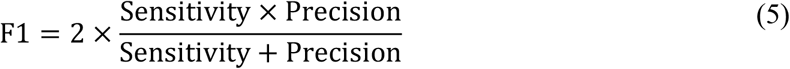

#### In trans variant identification

*In trans* variants are a key factor of PM3-relevant evidence. For our testing set, which contains 195 variant-publication pairs, with including total ground truth 294 *in trans* variants.

For evaluation, we have adopted a semi-automated way for evaluating the performance of *in trans* variant identification as the following procedure:

Given:

- Ground truth *in trans* variants *GT*
- Responses of the system *R*

1. For each ground truth *in trans* variant *g* ∈ *GT*:
  1.1. Generate all possible formats *F* for *g*, including DNA change, protein change, and homozygous state if applicable.
  1.2. Initialize flags *good*_*hit* and *suspect*_*hit* to empty lists.
  1.3. For each response *r* ∈ *R*:
    1.3.1. If any format *f* ∈ *F* is found in *r*, append *f* in *good*_*hit*.
    1.3.2. If *good*_*hit* is False and the position of any format *f* ∈ *F* matches the position in *r* within a tolerance, append f in *suspect*_*hit*.
  1.4. Save the *good*_*hit* and *suspect*_*hit*.
2. Manually review the responses with at least one *good*_*hit* or *suspect*_*hit*, and update the count of correctly identified *in trans* variants accordingly.

The recall was calculated based on the manually reviewed results as:

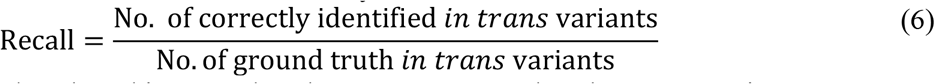

Noted that in our benchmarking samples, there are 294 ground truth *in trans* variants.

## Authors’ contributions

RL and SL contributed to the conceptualization and conceived the study. RL supervised the study. YW, CL, SL collected the data, implemented the algorithm, performed benchmarking evaluation. SL, CL, YW, RL, YH participated in manuscript drafting. All authors have participated in the result analysis, read and approved the manuscript.

## Code and data availability

Source code of AutoPM3 and PM3-Bench are accessible at: https://github.com/HKU-BAL/AutoPM3. The GGUF format of fine-tuned LLaMA3 model is available at: http://bio8.cs.hku.hk/AutoPM3. The online web server is accessible at: https://www.bio8.cs.hku.hk/autopm3-demo/.

## Acknowledgement

R.L. was supported by Hong Kong Research Grants Council grants GRF (17113721) and TRS (T21-705/20-N), the Shenzhen Municipal Government General Program (JCYJ20210324134405015), and the URC fund from HKU.

