## Supplementary Materials for "AutoPM3: Enhancing Variant Interpretation via LLM-driven PM3 Evidence Extraction from Scientific Literature"

### Supplementary Methods

#### Prompts and predefined queries of RAG model

The prompt of RAG module for both variant hit and *in trans* variant identification:

```
<|begin_of_text|><|start_header_id|>system<|end_header_id|>
You are a specialist in biogenetics, answer only based on user's input!<|eot_id|>
<|start_header_id|>user<|end_header_id|>
The variant in HGVS format is {question}, don't include this in your answer if condisering compound het variants.
Given the context: '{context}' and target variant {c_variant}. Answer the question: {proposedQuestion}<|eot_id|>.
<|start_header_id|>assistant<|end_header_id|>
```

The predefined query for the task of variant hit:

```
Does the paper mention the queried variant ({current_variant}) and what is the surrounding context? if such variant is existed, say *YES* at first otherwise say *None* (focus on variant: {current_variant})
```

The predefined query for the task of *in trans* variant identification:

```
If {current_variant} is compound heterozygous with another variant, name it; if {current_variant} is homozygous, say homozygous; if no related variant is found, say *None*. List all results separated by comma.
```

#### Prompts and predefined queries of TableLLM

The prompt of TableLLM for generating the SQL commands:

```
Given an input question, first create a syntactically correct {dialect} query to run, then look at the results \
of the query and return the answer. Strict your query to a short one and dont give a long answer. *Never* use limitation to limit your query like: LIMIT {top_k} except user asked for certain row.\nWhen no specific column names are given, you can check for the answer in all columns using "OR" operator.\n\n\
Unless exactly match is required by user, use LIKE other than = in the query\n\
Never sort the results. If user asks for certain row, use LIMIT operator!\n\
Never give a sql that will return all content in the table if not explicitly asked\n\
Only give one query ended with \';\' eachtime!\n\
```

```
Carefully check the statement after WHERE clause, don\'t mix up column_name with
user\'s query string, and keep the string integral for matching!\n\
When using LIKE operator, note to put column names on the left and query string on the
right, don\'t reverse it\n\
Don\'t forget to append ; at the end of query and no order is needed!\n\nPay attention to
use only the column \
names that you can see in the schema description. Be careful to not query for columns
that do not exist. Also, pay \
attention to which column is in which table.\n\nUse the following format:\n\nQuestion:
Question here\nSQLQuery: SQL \
Query to run\nSQLResult: Result of the SQLQuery\nAnswer: Final answer here\n\nOnly
use the \
following tables:\n{table_info}\n\nQuestion: {input}

get the first row in table {current_table}? (take the result given by SQLResult:)"

search for the string: '{query_variant}' through every column in table {current_table} using
OR? (find all, no limit, column names should be like 0,1,2 as u can see in the schema)"
```

The predefined queries:

```
find all rows that contain the string '{query_variant}' in any column (don\'t only consider
one column) (check all columns in table {current_table}) (find all, no limit)"
```

```
find all the rows that contain {query_variant} (query all columns in table {current_table}
using OR) (find all, no limit)
```

The prompt of TableLLM for convert the database query results into texts:

```
### System:
You are reading the structured data given in the Context and try to rephrase it in plain text. In each line, the attribute name(header) is on the left of *:*, then corresponding attribute value is on the right.

### Context:
{tableData}

### User:
Each variant/mutation must contain alphabet letters with several digits, don't make up non-existed variants/mutations.
Limit your answer under 25 words.
Stop the answer by the word *END*.
Please read the above provided structured data in context and just answer the given question in short plain text. Question: {question}'\
### Response:
```

Predefined query for summarization of the fetched results:

```
rephrase and describe it in plain text
```

Predefined query for extracting variant and patient information from the fetched results:

```
only list the existed variants/mutations in context in the following format *PatientID:... Variant:...*\nif no patient is explicitly mentioned put *PatientID:None* and don't mix up with variants/mutations. If no variants/mutations is explicitly mentioned put *Variant:None*
```

#### Fine-tuning data sample


```
{
 "conversations": [
  {
   "from": "user",
   "value": "Given the context: 'Two missense mutations causing mild HPA associated
with haplotype 12 88 swedish families. 93 children picked up by NBS A322G / R408Q –
300uM/L A322G / R408W – 250uM/L A322G / R408W – 360uM/L A322G / R252W –
250uM/L' and target variant in two format c.965C>G/p.(Ala322Gly). If target variant is
compound heterozygous with another variant, name it; if the target variant is
homozygous, say homozygous; if no related variant is found, say NA. List all results
seperated by comma."
  },
  {
   "from": "assistant",
   "value": "The variants contains:R408Q,R408W,R252W."
  }
 ],
 "id": 1
},
```

Supplementary Table 1. Variant hit performance of AutoPM3, vanillaRAG, and PaperQA.

| Methods | Sensitivity | Specificity | Precision |
|---|---|---|---|
| AutoPM3 | 0.892 | 0.83 | 0.839 |
| VanillaRAG (Llama3:8B) | 0.748 | 0.8 | 0.789 |
| VanillaRAG (Llama3:70B) | 0.558 | 0.953 | 0.922 |
| PaperQA (Llama3:70B) | 0.468 | 0.962 | 0.924 |

Supplementary Table 2. Variant hit performance of different models by sequentially add key AutoPM3's key modules

| Model | Sequentially added modules | Sensitivity | Specificity | Accuracy |
|---|---|---|---|---|
| Llama3:8B | Vanilla RAG | 0.748 | 0.8 | 0.774 |
| | Variant augmentation | 0.794 | 0.692 | 0.743 |
| | Variant retriever | 0.882 | 0.81 | 0.846 |
| | TableLLM | 0.815 | 0.876 | 0.845 |
| | Fine-tuning | 0.892 | 0.83 | 0.861 |
| Llama3: 70B | Vanilla RAG | 0.558 | 0.953 | 0.755 |
| | Variant augmentation | 0.62 | 0.953 | 0.786 |
| | Variant retriever | 0.876 | 0.989 | 0.983 |
| | TableLLM | 0.887 | 0.871 | 0.879 |
| Mistral-Large: 128B | Vanilla RAG | 0.205 | 0.984 | 0.594 |
| | Variant augmentation | 0.292 | 0.994 | 0.643 |
| | Variant retriever | 0.81 | 0.994 | 0.902 |
| | TableLLM | 0.887 | 0.876 | 0.881 |

Supplementary Table 3. *In trans* variant identification performance of different models by sequentially add key AutoPM3's key modules.

| Model | Sequentially added modules | Recall |
|---|---|---|
| Llama3:8B | Vanilla RAG | 0.092 |
| | Variant augmentation | 0.229 |
| | Variant retriever | 0.314 |
| | TableLLM | 0.709 |
| | Fine-tuning | 0.725 |
| Llama3: 70B | Vanilla RAG | 0.225 |
| | Variant augmentation | 0.306 |
| | Variant retriever | 0.540 |
| | TableLLM | 0.758 |
| Mistral-Large: 128B | Vanilla RAG | 0.197 |
| | Variant augmentation | 0.274 |
| | Variant retriever | 0.5 |
| | TableLLM | 0.745 |
